# Levamisole-Mediated Suppression of B-Cell Proliferation and Antibody Production Reveals Mechanistic Insights into Idiopathic Nephrotic Syndrome Therapy

**DOI:** 10.64898/2025.12.23.695947

**Authors:** Gerarda H. Khan, Floor Veltkamp, Nike Claessen, Loes Butter, Antonia H.M. Bouts, Sandrine Florquin, Jeroen E.J. Guikema

## Abstract

Levamisole (LMS) is a small imidazole derivative with immunomodulatory properties. Despite its longstanding clinical use in Idiopathic Nephrotic Syndrome (INS), the mechanisms underlying its immune-regulating effects remain poorly defined. In this study, we investigated the in *vitro* effects of LMS treatment on human B-cells, and in patients included from the LEARNS (**LE**vamisole as **A**djuvant therapy to **R**educe relapses of **N**ephrotic **S**yndrome) clinical trial. *In vitro* experiments showed that LMS directly suppresses activation and proliferation in both T-cell dependent and T-cell independent stimulated B-cells, without inducing cytotoxicity. In agreement, transcriptomic profiling demonstrated downregulation of cell cycle-associated genes and genes involved in immunoglobulin synthesis, while genes involved in terminal B-cell differentiation were upregulated, including SLAMF7. Consistently, LMS treatment decreased immunoglobulin expression and secretion while simultaneously inducing the expression of factors associated with a more terminal B-cell phenotype (CD138/CD319). Flowcytometry analysis of blood samples from LMS-treated INS patients revealed reduced circulating B-cell counts. Collectively, these data suggest that LMS acts as a potent modulator of B-cell function that inhibits proliferation and immunoglobulin synthesis, providing mechanistic support for its therapeutic efficacy in INS management.

## Introduction

Levamisole (LMS) is a small molecule and synthetic imidazole derivative known to inhibit tissue non-specific alkaline phosphatase (TNAP) [1] while also acting as an activator of nicotinic acetylcholine receptors [2]. It has long been recognized for its broad immunomodulatory properties [3] and has been used to treat a wide range of immune-related diseases [4–6]. Notably, LMS has proven effective in the treatment of idiopathic nephrotic syndrome (INS) [7, 8], a pediatric kidney disorder characterized by proteinuria, edema, and hypoalbuminemia.

INS is a complex disease in which injury of podocytes results in proteinuria [9]. Although the precise mechanism underlying podocyte injury in INS is unknown, beneficial clinical effects of the wide variety of immunosuppressive agents that are used to treat INS, suggest an immunological etiology [7]. Notable drugs used in the treatment of INS are corticosteroids, calcineurin inhibitors and (more recently) rituximab, which have all been shown to effectively induce or maintain remission in patients [7, 10].

While most INS patients initially respond well to steroids, approximately 70 percent will eventually relapse [11], and around 50 percent develop a frequent-relapsing pattern [12]. Frequent relapsers are at risk of developing steroid toxicity due to cumulative exposure associated with severe side effects. To prevent relapses and therefore steroid toxicity, additional immunosuppressive agents such as calcineurin inhibitors and rituximab are often introduced. However, both rituximab and calcineurin inhibitors are associated with their own severe side effects such as serious infections [13], seizures, hypertension and renal failure [14]. LMS has been associated with reducing the occurrence of relapses in frequently relapsing INS patients with relatively few side effects [15–18]. We are part of the LEARNS (**LE**vamisole as **A**djuvant therapy to **R**educe relapses of **N**ephrotic **S**yndrome) consortium, which is conducting a randomized clinical trial on the effect of LMS on relapse rates in INS and investigating the etiology of the disease [19].

Although T-cells have historically been implicated in the pathogenesis of INS [20–23], recent work highlights a potential pathogenic role for B-cells. A recent study by Hengel et al (2024) identified the presence of antibodies directed against nephrin in INS patients, an important protein in the cytoskeleton of podocytes [24]. Consistent with this data, flowcytometry analysis of INS patients with active disease has shown higher frequencies of total circulating B-cells, memory B-cells [25, 26] and plasmablasts [27].

The mechanism from which the immunomodulative effects of LMS originate is not yet understood. Several studies have reported its effects on T-cells [28–35]. Our group has previously shown that LMS suppresses T-cell activation and proliferation [36]. In contrast to the literature describing effects of LMS on human T-cells, B-cells have received relatively little attention. Prior studies have both described elevation of B-cell proliferation and immunoglobulin (Ig) production [37, 38] and reduced numbers of immunoglobulin-producing cells in treated volunteers [39], as well as a decrease of proliferation in human myeloma cell lines [40]. However, a detailed characterization of the mechanism of action of LMS in human B-cells is currently lacking.

In this study, we aimed to elucidate the effects of LMS on human B-cells. Here, we show that LMS directly suppresses B-cell activation and proliferation *in vitro*, both in T-cell mediated and T-cell independent stimulated B-cells. We further demonstrate that treatment with LMS induces cell cycle inhibition and reduces immunoglobulin production. In addition, we describe that LMS induces a phenotype associated with terminal B-cell differentiation. Lastly, we show that INS patients from the LEARNS trial treated with LMS display reduced circulating B-cell numbers, in accordance with its suppressive effects on B-cells *in vitro*. Together, these results indicate that LMS directly modulates human B-cell function and exerts potent immunosuppressive effects, supporting its therapeutic role in the management of INS.

## Materials and Methods

### Patient samples

Patient samples were gathered through the LEARNS study (**LE**vamisole as **A**djuvant therapy to **R**educe relapses of **N**ephrotic **S**yndrome) [19], an international double-blind placebo controlled randomized trial. Patients aged 2-16 years presenting with a first episode of INS were included in the trial and treated with oral prednisolone. After 4 weeks of successful treatment, patients were randomly allocated to additionally receive either LMS 2.5 mg/kg per alternating day or placebo. After 4 weeks, the prednisolone was slowly tapered till it was stopped at week 18 and patients only received study medication[19]. Patient blood samples were collected at week 20 and peripheral mononuclear blood cells (PBMCs) were isolated and analyzed through flow cytometry.

### Sex as a biological variable

In this study, sex was not considered as a biological variable.

### Isolation and in vitro culturing of primary human B-cells

Human PBMCs were isolated from six buffy coats using Ficoll-Hypaque density gradient centrifugation. The buffy coats were obtained from adult anonymous healthy blood donors (Sanquin Blood Supply, Amsterdam, The Netherlands), all of whom provided written informed consent for the use of residual blood in research. B-cells were isolated from PBMCs by negative magnetic-activated cell sorting (MACS) using the human pan B-cell isolation kit (Miltenyi Biotec, Bergisch Gladbach, Germany) following the manufacturer’s protocol. The purity of the B-cell fraction was verified by flow cytometry using CD19 (>92% purity for all donors). Purified B-cells were counted with a Bürker-Türk counting chamber and cryopreserved in fetal calf serum (HyClone, GE Healthcare Life Sciences, Chicago, IL, USA) supplemented with 10% dimethyl sulfoxide at a concentration of 10 million cells/mL. Cryovials were initially stored in a CoolCell freezing container (Corning, Sigma-Aldrich, St. Louis, MO, USA) at −80°C for 24 hours, after which they were transferred to liquid nitrogen. Prior to experiments, B-cells were thawed and rested overnight at a density of 2 million cells/mL in Iscove’s modified Dulbecco’s medium (IMDM; GIBCO, Paisley. U.K.) supplemented with 2 mM L-glutamine, 100 U/mL penicillin, 100 μg/mL streptomycin (Gibco, Thermo Fisher Scientific, Waltham, MA, USA), and 10% fetal calf serum. For experimental cultures, B-cells were maintained at a density of 1 × 10^6 cells/mL at 37°C with 5% CO2. At the start of activation, B-cell samples were treated with 500 µM or 10 µM Levamisole (levamisole-hydrochloride; Sigma-Aldrich), or 1:200 µL H2O (solvent control condition). LMS was refreshed after 24 h.

### Proliferation experiments

B-cells were activated through two distinct methods: T-cell dependent and T-cell independent activation. T-cell dependent activation was achieved by co culturing with plate bound lethally irradiated murine 3T3 cells expressing CD40L (3T3-CD40L) in the presence of recombinant human IL-2 (Peprotech, Thermo Fisher Scientific, Waltham, MA, USA), IL-4 and IL-21(Prospec protein specialists, Hamada, Israel). For T-cell dependent activation experiments, 20.000 lethally irradiated 3T3-CD40L cells per well were seeded into flat-bottom 96-wells plates in complete IMDM, supplemented with IL-2 (50 µg/mL), IL-4 (50 ng/mL) and IL-21 (50 ng/mL). The co-culture was maintained for 48 - 96 h. In some experiments, B-cells were collected from the plate-bound 3T3-CD40L cells and cultured for an additional 48 hours in complete IMDM supplemented with IL-2, IL-4 and IL-21 in absence of 3T3-CD40L.

T-cell independent activation was achieved through culturing with the human TLR9 ligand class B CpG oligonucleotides (ODN 2006, Invivogen, Toulouse, France). B-cells were cultured for 72 - 120 h in complete IMDM supplemented with CpG oligonucleotides (5 µg/mL).

After activation, supernatants were removed and the cells were collected or stained for further analysis.

### Flow cytometric analyses

Activated B-cells were washed with FACS buffer (0.2% BSA, 0.5 mM EDTA, 0.1% sodium azide in PBS) and incubated with the respective antibodies/dyes for 30 minutes on ice, protecting them from light. Flow cytometric analysis was performed using a BD LSRFortessa, and data were analyzed with FlowJo software (v10.5; FlowJo LLC, Ashland, OR, USA). Gates were set based on Fluorescence Minus One (FMO) staining, and compensation was performed using single-stain controls.

The following antibodies/dyes were used: CD19-PC7 (J3-119; Beckman Coulter; Brea, CA, USA), CD20-Pacific Blue (2H7; Biolegend; San Diego; CA; USA), CD38-APC-H7 (HB7; Becton Dickinson; Franklin Lakes; NJ; USA), CD69-PE (L78; Becton Dickinson), CD138-PE (S22003D; Biolegend), CD319-APC (CRACC; Biolegend), IgM-PE (MHM-88; Biolegend), IgG-PE-Cy7 (M1310G0S; Biolegend), CellTrace CFSE (Invitrogen; Thermo Fisher Scientific), 7-aminoactinomycin-D ( BioLegend).

For cell cycle analysis, BrdU incorporation was used as described previously [41]. Briefly, cells were incubated for 1 hour with 40 μM BrdU (Thermo Fisher Scientific), fixed in 75% cold ethanol, and digested with pepsin and HCl. BrdU incorporation was assessed by staining with anti-BrdU-FITC (#347583; Becton Dickinson) and TO-PRO-3 (Thermo Fisher Scientific), followed by flow cytometric analysis.

### Immunoglobulin production

IgM and total IgG levels in supernatants were measured by using Enzyme-Linked Immunosorbent Assay (ELISA) as previously described [42]. In short ELISA plates (Half area high binding 96-wells plates, Greiner bio-one, Kremsmünster, Austria) were coated overnight with monoclonal antibodies (MAbs), respectively CLB-MH15/1 (anti-IgM) and CLB-MH16/1 (anti-IgG) at a concentration of 4 µg/mL in PBS. Plates were then subsequently washed with PBS/0,02% Tween, blocked with PBS/1% BSA, washed again and incubated with sample dilutions (1:1 in PBS) for 1 h. The calibration curve was plated by using serial dilutions of a standard serum (CLB-H003). The plates were washed and incubated with the respective MAbs conjugated to Horseradish peroxidase (HRP) for 2 h. After washing, the plates were treated with a substrate solution consisting of Tetramethylbenzidine 100 µg/mL, 0,003% H_2_O_2_ in 0,1 M sodium acetate buffer. The subsequent color reaction was stopped by adding 4 M H2SO_4_. Extinction was measured at 450 nm by using a clariostar plate reader. Ig concentrations were calculated based on the calibration curve.

### Gene expression profiling by RNA sequencing

RNA sequencing was performed on activated human peripheral blood B-cells obtained from three healthy donors. Total B-cells (CD19^+^) were activated for 48 h with irradiated 3T3-CD40L cells and IL-2, IL-4 and IL-2, treated with 500 µM LMS or solvent (untreated controls). RNA was isolated using TRIzol (Thermo Fisher Scientific) and the RNeasy Mini Kit (Qiagen, Hilden, Germany)), following the manufacturer’s guidelines. RNA quality was assessed with the RNA integrity number (RIN) on an Agilent TapeStation system, with all RNA samples showing a RIN >7.

Sequencing libraries were prepared using the TruSeq Stranded mRNA Library Kit and sequenced on the NovaSeq platform. Quality control, trimming, alignment, and quantification were performed using the Galaxy web platform (https://usegalaxy.org/) [43]. All reads showed a PHRED score >28. The resulting normalized feature counts table was both uploaded to and analyzed by using R2: Genomics Analysis and Visualization Platform (http://r2.amc.nl) and analyzed in R studio. Differential expression was analyzed using DESeq2 [44]. The false discovery rate was controlled for using the Benjamini–Hochberg correction, and genes with an adjusted p-value < 0.05 and a log2 fold change (Log2FC) > 0.5 were considered differentially expressed. Enrichment plots and leading edge analysis were performed by using the Gene Set Enrichment Analysis software (GSEA, v4.4.0, https://www.gsea-msigdb.org/)[45]

### Statistics

Statistical analyses were conducted using GraphPad Prism software (version 8.0.0 for Windows). Median fluorescence indices and geometric means of fluorescence were calculated using FlowJo software (version 10). Proliferation analysis was performed using the FlowJo Proliferation module, which quantifies cell divisions based on CFSE peaks. The division index was calculated by dividing the total number of divisions by the calculated number of cells at the start of the culture.

### Study approval

Written informed consent was obtained from parents and/or patients. This study was approved by the Medical Ethical Committee (MEC) of Amsterdam UMC (NL61906.018.17, 16 nov 2017). This study started recruiting patients in April 2018 and is currently in follow up and blinded. Buffy coats from anonymous healthy blood donors were obtained from Sanquin Blood Supply, Amsterdam, The Netherlands. All donors provided written informed consent for usage of the remainder of their routine blood donation in research. All experiments were performed according to the ethical standards of the institutional medical ethical committee of the Amsterdam UMC, as well in agreement with the declaration of Helsinki as revised in 1983.

### Data availability statement

The data presented in this study are accessible in the Gene Expression Omnibus (GEO), accession number: GSE314752

## Results

### Levamisole is an inhibitor of B-cell proliferation and exerts its effects independently of mode of stimulation

To gain insights into the effects of LMS treatment in our pediatric patient cohort, peripheral blood samples were analyzed by flow cytometry. In patients receiving LMS treatment (2.5 mg/kg every two days), a significant reduction in circulating B-cell percentages was observed compared to placebo-treated controls (Figure 1A).

**Figure 1.**
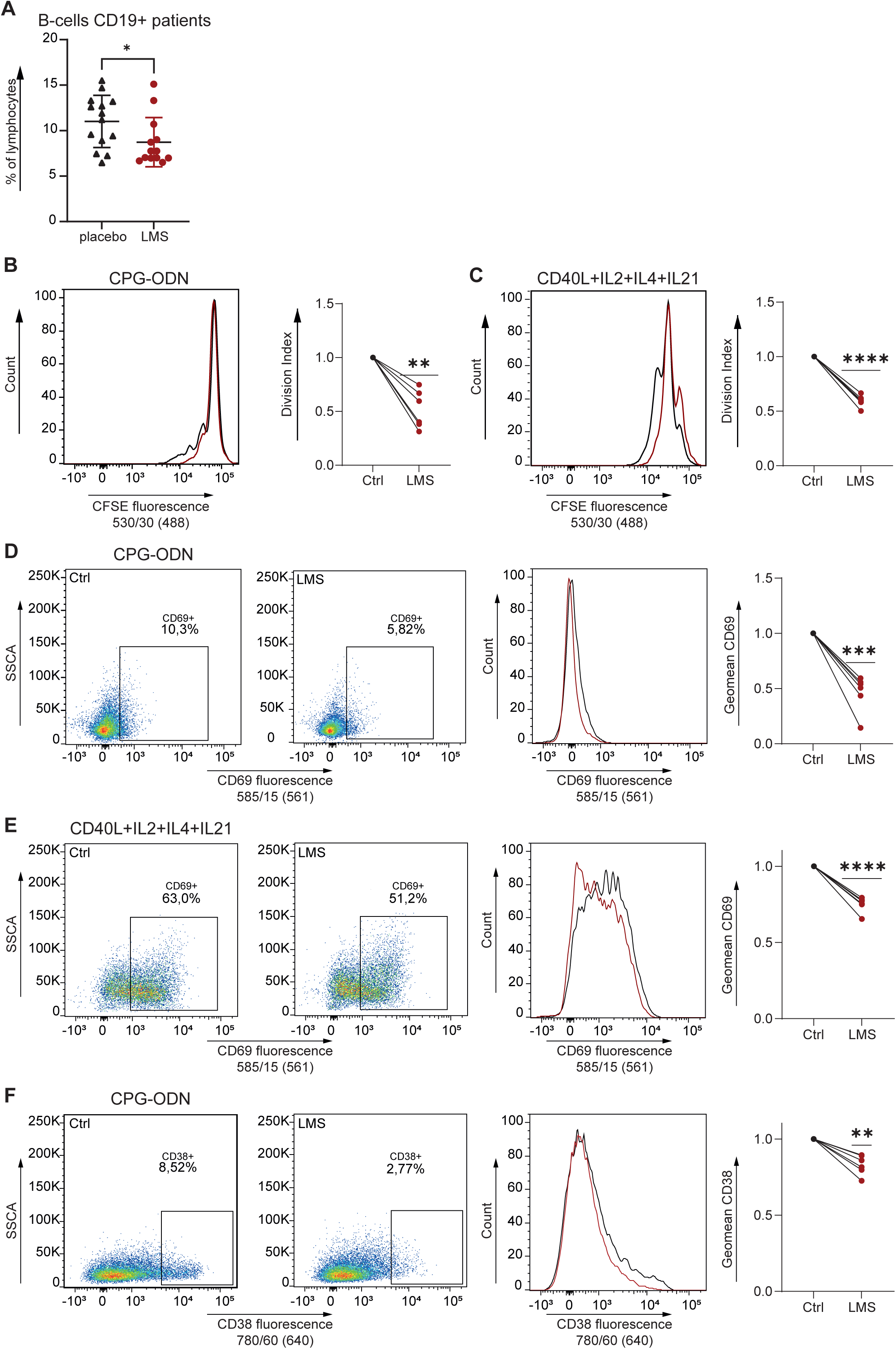
LMS decreases proliferation and activation in both T-cell dependent and independent activated human B-cells. (A) Graph depicting patients with INS from the LEARNS trial at week 20 in their treatment schedule. Patients receiving LMS (red, n =13 patients) show less circulating B-cells (CD19+) than patients receiving placebo (black, n = 14 patients) (*P ≤ 0.05, unpaired t-test). (B) LMS treatment decreased proliferation in T-cell independent activated B-cells as measured by the CFSE proliferation assay. B-cells (n = 6 donors) were stained with CFSE and subsequently activated by CpG (5 µg/mL) for 96 h. Samples were treated with 500 µM LMS (red) or solvent control (black) for the duration of activation. CFSE fluorescence was measured by flow cytometry (left) and quantified by gating the respective division peaks and calculating division indices. The uttermost right graph shows normalized division indices for 6 donors, the lines connect the same individual donor (** P ≤ 0.01, one sample t-test). (C) LMS treatment showed a similar decrease of proliferation in T-cell dependent activated B-cells as shown by CFSE fluorescence. CFSE stained B-cells (n = 6 donors) were activated with 3T3-CD40L cells 10:1 in combination with cytokines for 72 h. Fluorescence was measured (left) and quantified by calculating the division indices. The uttermost right graph shows normalized division indices for 6 donors (**** P ≤ 0.0001, one sample t-test). (D) LMS decreased the expression of CD69 expression in T-cell independent stimulated B-cells treated with LMS. B-cells (n = 6 donors) were stimulated with CpG for 96 h and treated with LMS 500 µM. Dot plots (left) and histograms (middle) for CD69 fluorescence are shown. CD69 fluorescence was quantified by calculating the fluorescence geometric means (right) for both LMS treatment (red) and solvent control (black). Normalized data are shown (***P ≤ 0.001, one sample t-test). (E) Similarly, LMS treatment decreased expression of CD69 expression in T-cell dependent stimulated B-cells. B-cells (n = 6 donors) were stimulated with 3T3-CD40L and cytokines for 48 h and treated with LMS 500 µM. Dot plots (left) histograms (middle) and quantified normalized geometric means (right) for CD69 fluorescence are shown (***P ≤ 0.001, one sample t-test). (F) Lastly, LMS treatment decreased CD38 expression in B-cells stimulated with CpG for 96 h as shown by dot plots (left), histograms (middle) and quantified normalized geometric means of fluorescence (right) (** P ≤ 0.01, one sample t-test).

To further characterize the effects of LMS on human B-cell proliferation, CFSE stained B-cells from healthy donors were stimulated in the presence of LMS. LMS was titrated to assess dose-dependent effects on proliferation. Concentrations ranging from 250 to 1000 µM markedly reduced B-cell proliferation, whereas lower concentrations showed no detectable effects (Supporting Information Fig. 1B-E). A concentration of 500 µM was selected for subsequent experiments, as it consistently produced a robust effect across all assays. Importantly, live/dead staining confirmed that LMS treatment did not induce cell death of proliferating B-cells (Supporting Information Fig. 1F-G).

At 500 µM, LMS significantly reduced proliferation in both T-cell-dependent (Figure 1C) and - independent (Figure 1B) B-cell stimulations, indicating that its antiproliferative effects are independent of the activation pathway. In addition to its antiproliferative activity, LMS treatment also led to a marked decrease in the expression of the activation markers CD69 [46] and CD38 [47] (Figure 1 D-F). Furthermore, LMS treatment resulted in a consistent decrease of basal respiration and maximal respiration in the oxygen consumption rate, while not affecting glycolysis, as measured by Seahorse assays (Supporting information Fig 2 A-C). Collectively, these results demonstrate that LMS directly impairs human B-cell proliferation and activation through a mechanism independent of the mode of stimulation, without inducing cytotoxic effects.

**Figure 2.**
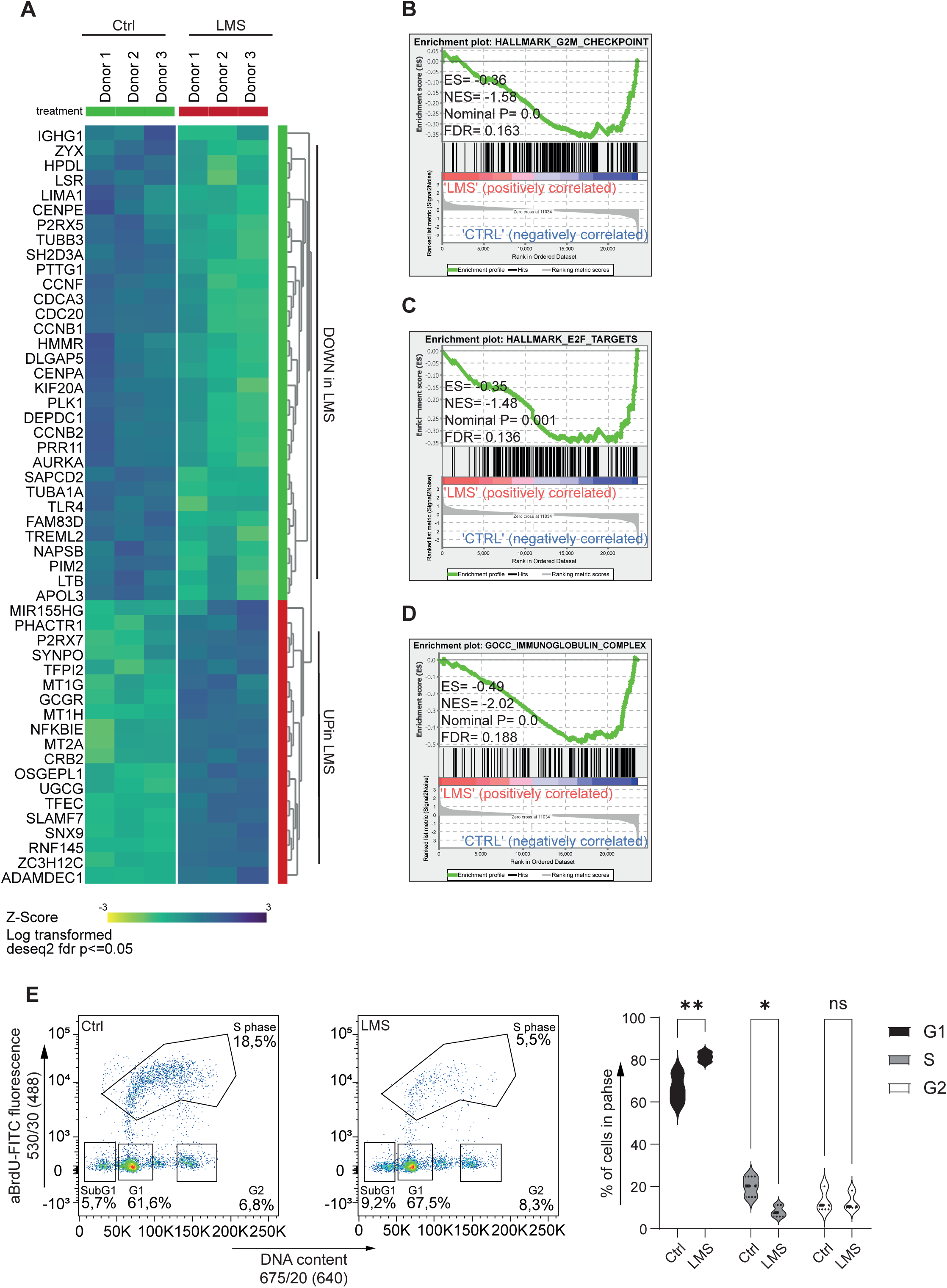
Gene expression profiling of LMS-treated T-cell depend activated B-cells shows downregulation of cell cycle associated genes and upregulation of genes associated with plasmablast differentiation. (A) Heatmap depicting differentially expressed genes in LMS-treated B-cells. In short B-cells were isolated from 3 donors and activated with 3T3-CD40L cells and cytokines for 48 h in combination with LMS 500 µM or solvent control. After activation, RNA isolation was performed and gene expression was examined through bulk RNA sequencing. Differentially expressed genes were defined as genes with an absolute Log2FC greater than 0.5 and with a p-value of <0.05. Heatmap shows log transformed z-scores. (B – D) Gene set enrichment analysis (GSEA) enrichment plots show negative enrichment of genes involved in cell cycle (B and C) and components of immunoglobulin synthesis (D). Shown are HALLMARK_G2M_CHECKPOINT (B), HALLMARK_E2F_TARGETS (C), GOCC_IMMUNOGLOBULIN_COMPLEX (D). The false discovery rate (FDR), normalized enrichment score (NES) and p-value are shown in the individual plots. (E) BrdU incorporation–based cell-cycle analysis showing the different phases: sub-G1 (dead cells), G1, S1 (early S phase), S2 (late S phase), and G2. Activated B-cells were treated with 500 µM or solvent control for 48 hours, after which BrdU incorporation and DNA content were assessed by flow cytometry. From left to right: dot plot graphs for cells treated with solvent control (left), LMS 500 µM (middle) and quantification by violin plots with comparison between the phases and treatment groups (G1 ** p < 0.01 and S * p < 0.05, two-way ANOVA with Šidáks multiple comparison test).

### Levamisole affects transcription of cell-cycle associated genes, immunoglobulin associated genes and genes associated with B-cell differentiation

To investigate the molecular mechanisms and potential targets of LMS, we performed RNA sequencing on activated human B-cells treated with LMS. Using a threshold of Log₂FC > 0.5 and p < 0.05, 51 genes were identified that were differentially expressed following 48 h of LMS treatment, including 32 downregulated and 19 upregulated genes. In line with the inhibitory effect of LMS on B-cell proliferation, several genes involved in cell-cycle regulation, such as *CCNB1, CCNB2,* and *PLK1*, were significantly downregulated (Figure 2A). This was further supported by gene set enrichment analysis which confirmed the depletion of gene sets involved in cell-cycle progression (Figure 2B-C) and cell-cycle analysis experiments which showed a significant inhibition in the progression from G1 to S-phase (Figure 2E).

Genes associated with immunoglobulin assembly, including *IGHG1* (Figure 2A) and *IGLV2* (Supporting Information Fig 3), showed a similar downregulation upon LMS exposure which was further supported by the depletion of genes involved in the assembly of the immunoglobulin complex by gene set enrichment analysis (Figure 2D).

**Figure 3.**
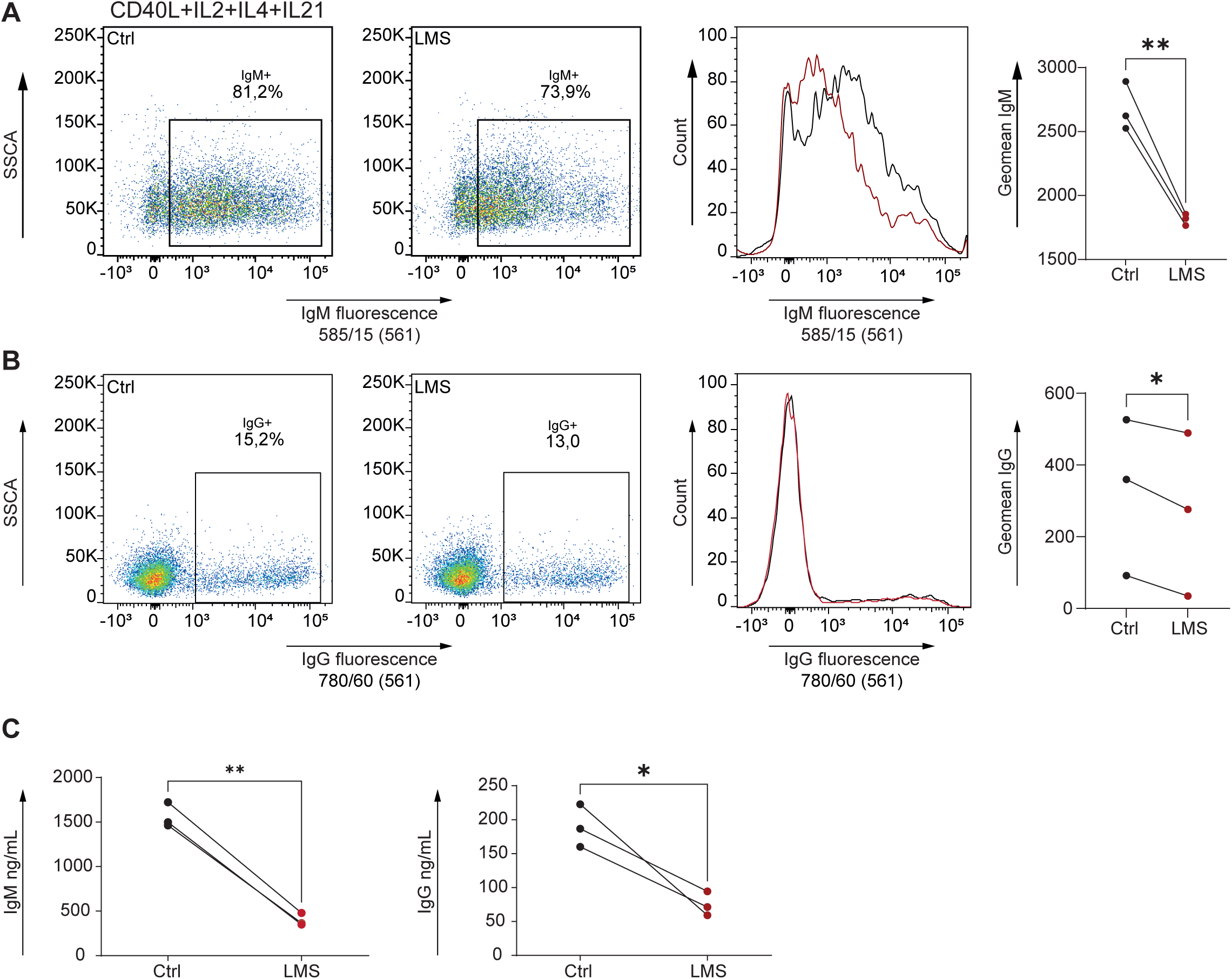
LMS treatment decreases IgM and IgG expression and secretion in activated B-cells. (A) plots depicting fluorescent expression of IgM by dot plots (left), histograms (middle) and quantification by geometric mean (right). B-cells were activated with 3T3-CD40L and cytokines for 48 h and treated with LMS 500 µM or solvent control. LMS treated B-cells are depicted in red (n = 3 donors, ** P ≤ 0.01, paired t-test). (B) plots depicting fluorescent expression of IgG by dot plots (left), histograms (middle) and quantification by geometric mean (right). B-cells were activated with 3T3-CD40L and cytokines for 96 h and treated with LMS 500 µM or solvent control (n = 3 donors, *P ≤ 0.05, paired t-test). (C) Graphs depicting IgM (left) and IgG (right) concentrations in supernatant after 96 h of activation by 3T3-CD40L and cytokines (n = 3 donors, **P ≤ 0.01, * p < 0.05, paired t-test).

Interestingly, LMS treatment induced an upregulation of *SLAMF7* transcription, which encodes CD319, a surface molecule strongly associated with differentiation towards memory B-cells and plasma cells, which implies that LMS could affect terminal differentiation in B-cells [48, 49].

Together, these findings suggest that LMS modulates B-cell activation by suppressing cell-cycle progression and immunoglobulin synthesis while also promoting transcriptional features of terminal differentiation.

### Levamisole reduces the expression and secretion of immunoglobulins

To further assess the impact of LMS on immunoglobulin production, we examined its effects on both the expression and secretion of IgM and IgG. LMS treatment markedly reduced IgM surface expression (Figure 3A) and caused a moderate reduction in IgG surface expression (Figure 3B) as measured by flowcytometry. Consistent with these findings, the secretion of both IgM and IgG was similarly diminished following LMS exposure (Figure 3C). These results indicate that LMS significantly impairs B-cell immunoglobulin production, reflecting its broader inhibitory effects on B-cell functional differentiation.

### Levamisole drives B-cell progression toward a terminal phenotype

The observed upregulation of *SLAMF7* transcription (Figure 2A) prompted us to further investigate the expression of markers associated with terminal B-cell differentiation. LMS treatment resulted in a significant decrease in CD20 expression after 48 hours (Figure 4A). In contrast, CD138 expression was significantly increased after stimulation and LMS treatment (Figure 4C). Both of these changes are indicative of a more plasmablast like phenotype [50]. Consistent with the *SLAMF7* transcription data, CD319 expression was markedly elevated after LMS treatment (Figure 4B).

**Figure 4.**
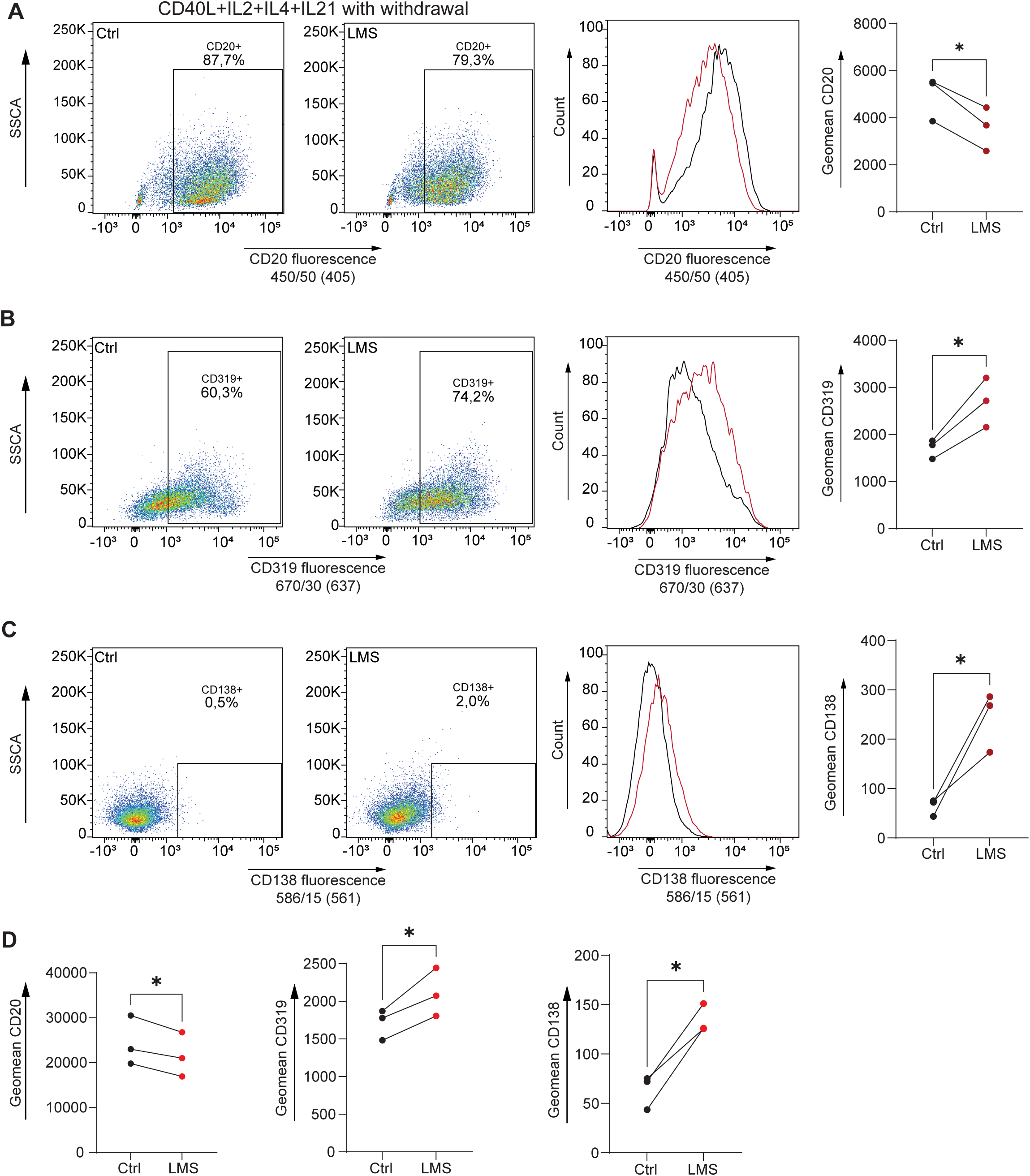
LMS treatment induces a more terminal B-cell phenotype with a decrease of CD20 and increase in CD319 and CD138 expression which is persistent at physiological concentrations. (A) graphs showing CD20 expression at 48 h, shown here in dot plots (left), histograms (middle) and quantified by geometric means (right) (*P ≤ 0.05, paired t-test). (B) and (C) showing similar graphs for CD319 at 72 h (B) and CD138 at 48 h activation and subsequent 24 hour rest without activation (C) with dot plots (left), histograms (middle) and quantified geometric means (right) (*P ≤ 0.05, paired t-test). All samples were activated with 3T3-CD40L cells and cytokines as specified by the specific timepoints and treated with LMS 500 µM (n = 3 donors). (D) Graphs depicting quantified geometric means for CD20 (left), CD319 (middle) and CD138 (right) expression for B-cells treated with LMS 10 µM (bright red) or solvent control (black). CD20 and CD138 expression was measured after 48 h activation and subsequent 24 h withdrawal of activation. CD319 expression was measured after 72 h. (n = 3 donors, *P ≤ 0.05, paired t-test).

Interestingly, while lower concentrations of LMS did not affect proliferation (Supporting Information Fig. 1B-E), similar effects on B-cell differentiation were observed at 10 µM (Figure 4D). At this concentration, LMS significantly reduced CD20 expression and increased both CD319 and CD138 expression.

## Discussion

As B-cells seem to play a role in the etiology of INS [25, 26] and therapeutic agents targeting B-cells have been proven effective in inducing remission in INS [7, 10], we aimed to examine the effects of LMS on human B-cells. Here, we show that INS patients treated with LMS exhibit decreased circulating B-cell (CD19+) numbers as compared to patients receiving placebo. We further show that *in vitro* treatment with LMS directly affects human B-cells and limits proliferation, activation and attenuates immunoglobulin production, independently of the mode of activation. Transcriptomic analysis indicated that LMS functions as an inhibitor of cell-cycle progression by limiting the cells to progress from G1 to S-phase. Simultaneously, LMS appears to promote features of terminal B-cell differentiation, as evidenced by the increase in expression of CD319, CD138 and decrease in CD20 expression. Together these findings show that LMS acts as a potent suppressor of B-cells and thereby may reduce the immunological drivers of disease activity in INS.

INS has long been considered a T-cell-driven disease [20–23], but increasing evidence implicates B-cells and humoral immunity in its pathogenesis [51]. The effectiveness of rituximab in maintaining remission in INS was a first crucial hint at the possible role for B-cells in its etiology [7, 10]. Studies focusing on the composition of B-cell populations in INS patients have shown conflicting results, with some studies reporting elevated B-cell numbers in active INS [25, 52], while other studies have not [53–56]. Among B-cell subsets, memory B-cells and short lived antibody secreting cells (plasmablasts) appear to be the most consistently elevated in INS patients [27, 52, 56, 57]. A recent study performed by Al-Aubodah et al showed an increase of marginal-zone like B-cells, atypical B-cells and antibody secreting cells in active INS patients, suggesting an aberrant extra follicular B-cell expansion associated with the formation of short lived antibody secreting cells [58]. Additionally, elevated IgM and decreased IgG levels have been previously documented in INS patients[59, 60]. In line with all these findings, low levels of circulating auto-antibodies directed at nephrin have been detected in a substantial portion of INS patients. In these studies, levels correlated to disease activity and depletion of these antibodies was observed after treatment with rituximab [24]. These results suggest that B-cells play an important role in the development of INS. Collectively, these results support a model in which earlier immune activation drives the formation of antibody-secreting cells and the production of autoreactive antibodies that target podocytes, leading to foot process injury. Because LMS is shown here to strongly inhibit B-cell activation and proliferation, it may be able to disrupt this pathogenic route.

Previous work from our group has demonstrated that LMS dampens the activation, proliferation, and cytokine production of T-cells by inducing cell cycle inhibition through a p53-dependent DNA damage response [36]. Here, we show that LMS inhibits B-cell activation, proliferation and immunoglobulin secretion by inducing cell-cycle arrest without transcriptional evidence of DNA damage or p53-upregulation (Supporting Information Fig 3 B-C), suggesting a different underlying immunosuppressive mechanism of LMS in B-cells. Furthermore, we show that human B-cells are uniquely sensitive to LMS, as shown by the lower concentrations needed to induce a robust immunosuppressive response (Supporting Information Fig. 1C and E), further supporting differences in reactivity to LMS between human B-cells and T-cells. We also show that LMS inhibits B-cell proliferation in both T-cell dependent and T-cell independent activated B-cells, indicating that LMS acts downstream in the activation pathway of B-cells. The reduction in proliferation we see during LMS treatment without inducing cytotoxic effects (Supporting Information Fig 1 F-G) indicates a cytostatic mechanism over a cytotoxic mechanism. This is further supported by the observed inhibition in cell-cycle (Figure 2 A-E) and the reduction in mitochondrial respiration (Supporting Information Fig. 2 A and C) in LMS-treated B-cells.

LMS acts as an inhibitor of Tissue-Nonspecific Alkaline Phosphatase (TNAP) [1, 30, 61] and an agonist of the nicotinic Acetylcholine Receptor (nAChR)[2, 62]. However, it remains unclear whether its immunomodulatory effects are mediated through these pathways. TNAP and its corresponding gene *ALPL* are expressed in both human and murine B-cells, and increased *ALPL* expression correlates with enhanced TNAP activity in murine B-cells[63]. In these murine B-cells, TNAP activity increases during B-cell proliferation[64], peaks during S-phase [65], and has been linked to differentiation into antibody secreting cells [64, 66]. Our experiments revealed no differences in the transcription of *ALPL* (Supporting Information Fig. 4B) indicating an alternate mechanism may be involved. Acetylcholine receptors are expressed on both murine and human B-cells [67]. nAChR activation in murine B-cells was shown to either promote or inhibit proliferation and IgG expression depending on the targeted subunit, specifically the activation of α7 of nAChR was associated with a decrease in B-cell proliferation [68–70]. LMS exhibits weak agonistic activity on levamisole-sensitive nAChRs in nematodes [71] and on α3β2 and α3β4 nAChR subunits in human neuronal cells [2].Whether LMS can activate the α and β subunits of nAChR found on human B-cells remains unknown. A recent study using an adjuvant-induced arthritis model in rats showed that LMS inhibits the PI3K/Akt/mTOR pathway [72], a key regulator of cell cycle progression and regulation, yet our experiments showed no reduction in its transcriptional activity (Supporting information Fig 4 C). Similarly, LMS reduced JAK/STAT signaling in both transcriptional and functional data [35], which we did not observe in our transcriptional data (Supporting information Fig 4 D), making this a less likely mechanism as well.

Interestingly, while LMS treatment decreased both proliferation, CD38 expression and Ig secretion in our experiments, it simultaneously increased the expression of several markers associated with terminal B-cell differentiation. Typically, the differentiation of quiescent B-cells into antibody secreting cells is preceded by extensive proliferation, an increase in CD38 and Ig expression and a decline in CD20 expression. Terminally differentiated antibody secreting cells characteristically express CD138 and CD319. During this process, B-cells usually upregulate *PRDM1*, *XBP1* and *IRF4* while the transcription of *PAX5* and *BACH2*, associated with a more naïve B-cell program, is suppressed [73]. Our data reveals atypical features of B-cell differentiation, not previously described in the context of LMS-treatment. Specifically, LMS upregulated CD138 and CD319 expression, even at low concentrations, despite its strong inhibitory effects on proliferation and Ig synthesis. At the transcriptional level we see a trend towards increase in the expression of *IRF4* (Supporting Information Fig 4 E). However, the transcription of other differentiation associated genes were not affected, and gene sets associated with terminal B-cell differentiation were not significantly enriched (Supporting Information Fig 4 A). A possible explanation for this seemingly contradictory finding is that LMS robustly suppresses B-cell proliferation. This may push B-cells into a cellular state best described as an “identity crisis” in which they display plasma cell-like characteristics while remaining in an arrested proliferative state-a phenotype previously linked to impaired cell survival [74].

This study demonstrates that LMS is a potent immunosuppressant of human B-cell activation and proliferation which could explain its therapeutic benefits in INS, a potentially B-cell driven disease. Our previous work has shown that LMS also suppresses T-cell activation and proliferation, which could indirectly decrease B-cell proliferation and differentiation through the suppression of CD4+ T helper cells. LMS is recognized as an immunomodulative drug with relatively mild side effects[7] and may therefore represent a promising adjuvant immunosuppressive agent in the management of INS. Although the exact mechanism of action of LMS in INS patients remains elusive, our group is currently investigating its immunomodulatory effects in this patient population. Ongoing and future research will provide further insight into the underlying mechanisms of LMS-mediated immune regulation.

## Author contributions

G.H.K., J.E.J.G. and S.F. designed the study and wrote the experiments. Experiments and analyses were performed by G.H.K., N.C. and L.B.. G.H.K. performed analyses on the sequencing dataset. A.H.M.B. is principle investigator of the LEARNS consortium and responsible for the study and collection of patient samples. F.V. and A.H.M.B. provided technical and theoretical input. Supervision of the study was given by J.E.J.G and S.F.. All authors read and approved the final manuscript.

## Funding support

This work was funded by the Dutch Kidney Foundation (DKF, CP 16.03) and the Rare Diseases Fund (Stichting Zeldzame Ziekten Fonds, the Netherlands).

## Acknowledgements

The authors would like to thank Berend Hooibrink, Kim Brandwijk-Paarlberg, and Toni van Capel for their suggestions and outstanding flow cytometry support.

## Notes

### Competing Interest Statement

The authors have declared no competing interest.

