## Supplemental information for "Levamisole-Mediated Suppression of B-Cell Proliferation and Antibody Production Reveals Mechanistic Insights into Idiopathic Nephrotic Syndrome Therapy"

### Supporting methods

**Metabolic profiling with Seahorse:**
Seahorse XF metabolic profiling to determine the mitochondrial metabolic function (XF cell Mito Stress Test) was carried out as per the manufacturer's instructions. In short: XF culture plates were coated with poly-D-lysin (50 µg/ml, Sigma Aldrich) and incubated overnight. Human B-cells were stimulated with 3T3-CD40L and a cytokine cocktail as described under material and methods for 48 h with LMS 500 µM or solvent control. After 48 h the cells were spun down and counted. 6 replicate measurements of 100.000 cells per replicate were used and taken up in 50 µL base medium (Dulbecco's modified eagle medium (DMEM), 45 µM phenol red, 31.6 mM NaCl, 25 mM D-glucose, 2 mM glutamin and 1 mM sodium pyruvate, pH 7.4) and plated in the XF plate. 125 µL base medium was added to each well and the plate was incubated for 1 h at 37°C in absence of CO2. The plate was then measured on a seahorse 96XFe analyzer (Agilent). During the assay, the following compounds were sequentially injected to reach final concentrations of 1.5 µM oligomycin (A), 2 µM FCCP (B), and 1.25 µM rotenone plus 2.5 µM antimycin A (C).

The following values were calculated:

Non mitochondrial respiration: the minimum rate measurement after last injection with Rotenone and antimycin A.

Basal respiration: the last rate measurement before the first injection with oligomycin – non mitochondrial respiration

Maximal respiration: maximum measurement after injection with FCCP – non mitochondrial respiration

Proton leak: minimum measurement after oligomycin injection – non mitochondrial respiration

ATP production: last measurement before injection with oligomycin – minimum measurement after oligomycin injection

Spare respiratory capacity: maximal respiration – minimum measurement after oligomycin injection.

### Supplemental 1

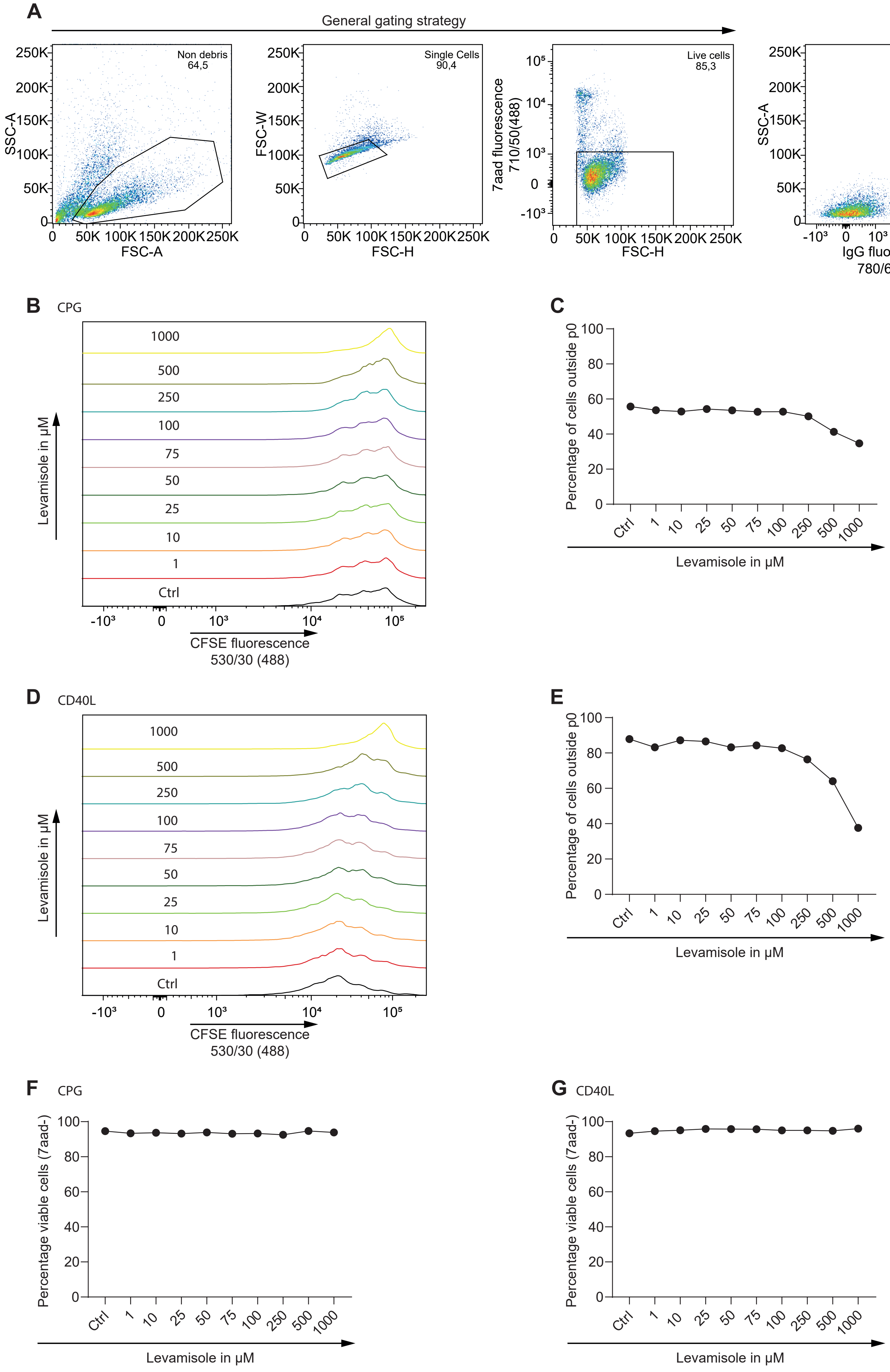

### Supplemental 2

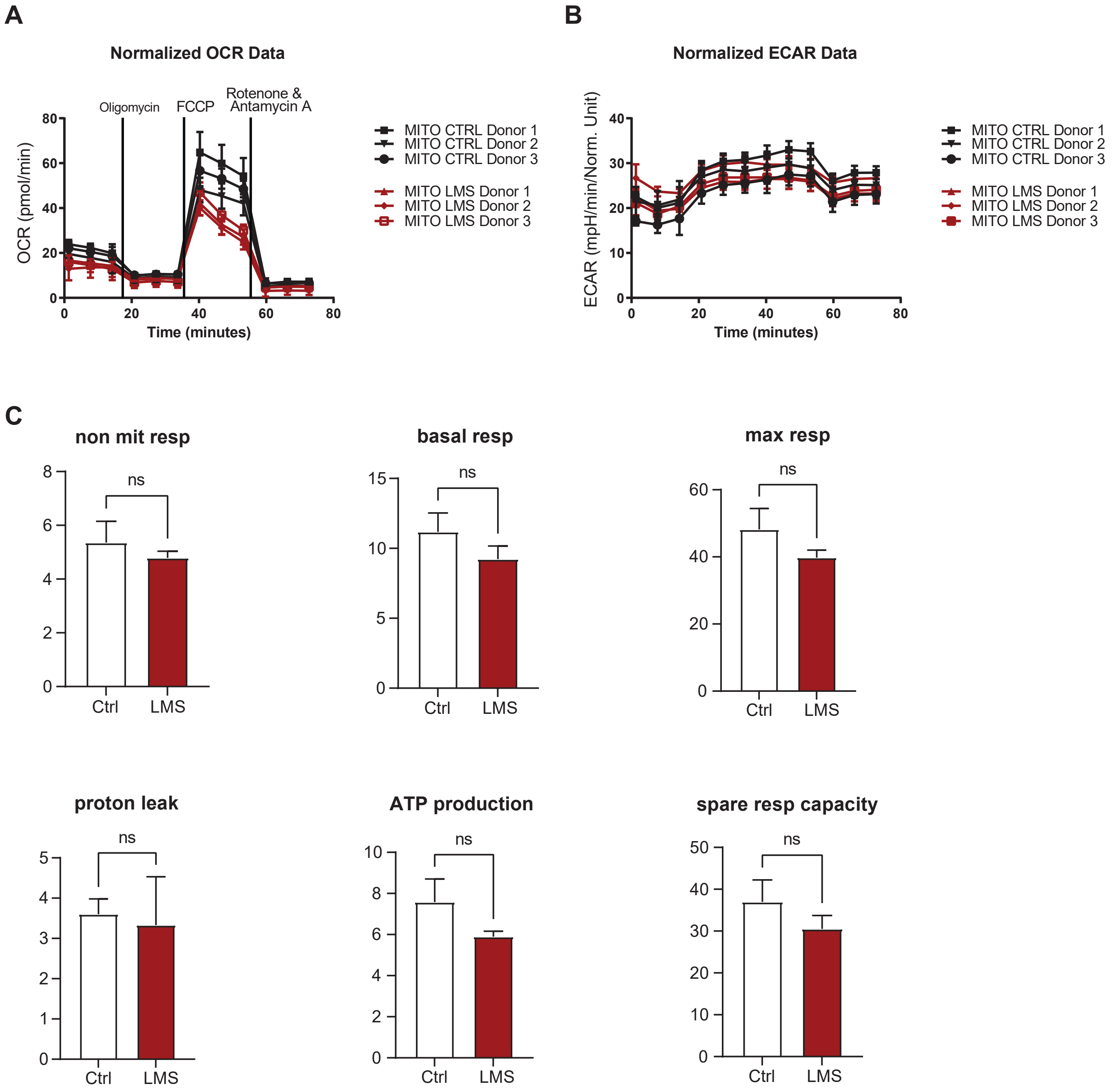

### Supplemental 3

## A

| HALLMARK_G2M_ CHECKPOINT_LE | HALLMARK_E2F_ TARGETS_LE | GOCC_IMMUNOGLOBULIN _COMPLEX_signal_LE |
|---|---|---|
| AURKA | ASF1A | CD79A |
| AURKB | AURKA | CD79B |
| BIRC5 | AURKB | IGHD1-1 |
| BUB1 | BIRC5 | IGHG1 |
| CCNA2 | BUB1B | IGHG2 |
| CCNB2 | CCNB2 | IGHG4 |
| CCNF | CDC20 | IGHM |
| CDC20 | CDC25B | IGHV1-18 |
| CDC25B | CDCA3 | IGHV1-45 |
| CDK1 | CDCA8 | IGHV1-69 |
| CDKN3 | CDK1 | IGHV2-26 |
| CENPA | CDKN3 | IGHV2-5 |
| CENPE | CENPE | IGHV2-70 |
| CENPF | CKS2 | IGHV3-13 |
| CKS2 | DCTPP1 | IGHV3-15 |
| DBF4 | DDX39A | IGHV3-20 |
| DTYMK | DEPDC1 | IGHV3-23 |
| E2F2 | DLGAP5 | IGHV3-30 |
| EFNA5 | E2F8 | IGHV3-33 |
| EWSR1 | GINS4 | IGHV3-35 |
| FBXO5 | HMGA1 | IGHV3-38 |
| HMGB3 | HMGB2 | IGHV3-43 |
| HMMR | HMGB3 | IGHV3-48 |
| HOXC10 | HMMR | IGHV3-7 |
| INCENP | ING3 | IGHV4-28 |
| JPT1 | JPT1 | IGHV4-4 |
| KIF23 | KIF18B | IGHV4-59 |
| KIF2C | KIF22 | IGHV5-51 |
| KIF4A | KIF2C | IGKC |
| KMT5A | KIF4A | IGKV1-13 |
| KPNA2 | KPNA2 | IGKV1-16 |
| LBR | LBR | IGKV1-27 |
| MEIS1 | MAD2L1 | IGKV1-37 |
| MKI67 | MCM5 | IGKV1-5 |
| NDC80 | MKI67 | IGKV1-6 |
| NEK2 | MXD3 | IGKV1-8 |
| NUSAP1 | MYBL2 | IGKV1D-17 |
| PBK | NAA38 | IGKV1D-39 |
| PLK1 | NCAPD2 | IGKV1D-43 |
| PLK4 | NME1 | IGKV2-24 |
| PRC1 | ORC2 | IGKV2-28 |
| PTTG1 | PA2G4 | IGKV2-29 |
| RACGAP1 | PCNA | IGKV2-30 |
| RBM14 | PLK1 | IGKV2-40 |
| SLC7A5 | PLK4 | IGKV2D-24 |
| TACC3 | POLD1 | IGKV3-20 |
| TLE3 | POP7 | IGKV3-7 |
| TOP2A | PTTG1 | IGKV3D-11 |
| TPX2 | RACGAP1 | IGKV3D-20 |
| TROAP | RANBP1 | IGKV4-1 |
| TTK | RNASEH2A | IGKV5-2 |
| UBE2C | RRM2 | IGLC2 |
| UBE2S | SMC4 | IGLL5 |
| | SNRPB | IGLV10-54 |
| | SPAG5 | IGLV1-36 |
| | SPC24 | IGLV1-40 |
| | SSRP1 | IGLV1-44 |
| | TACC3 | IGLV2-11 |
| | TBRG4 | IGLV2-14 |
| | TK1 | IGLV2-18 |
| | TOP2A | IGLV2-23 |
| | TP53 | IGLV2-8 |
| | TRIP13 | IGLV3-1 |
| | TUBB | IGLV3-10 |
| | UBE2S | IGLV3-19 |
| | | IGLV3-21 |
| | | IGLV3-25 |
| | | IGLV3-27 |
| | | IGLV3-9 |
| | | IGLV4-3 |
| | | IGLV7-46 |
| | | IGLV8-61 |
| | | JCHAIN |
| | | LIME1 |
| | | VPREB3 |

## B

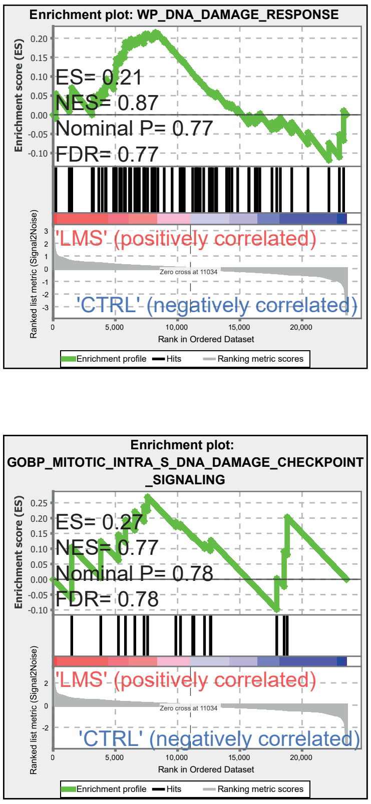


## C

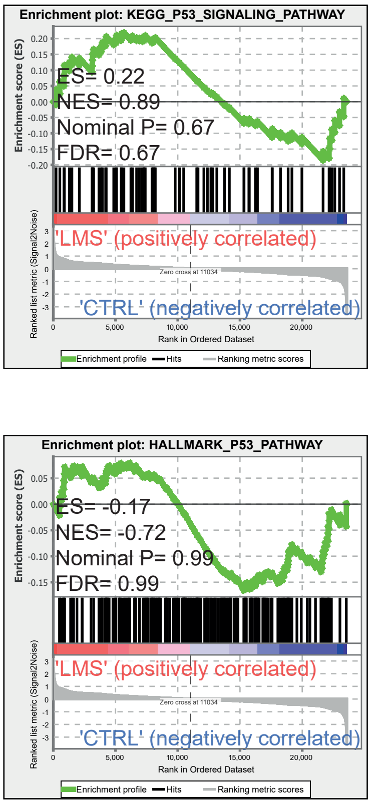

### Supplemental 4

A

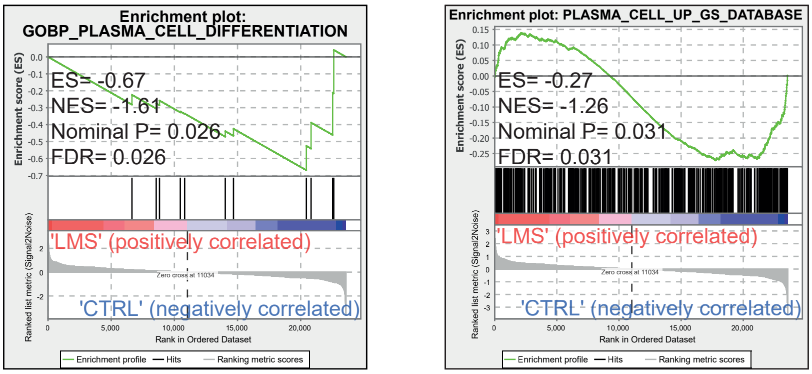


B

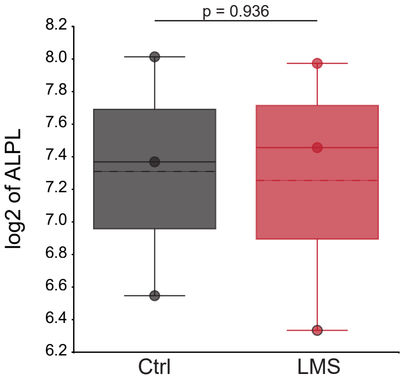


C

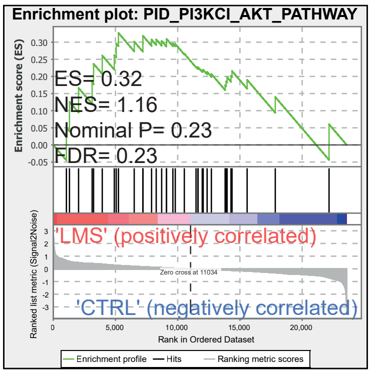


D

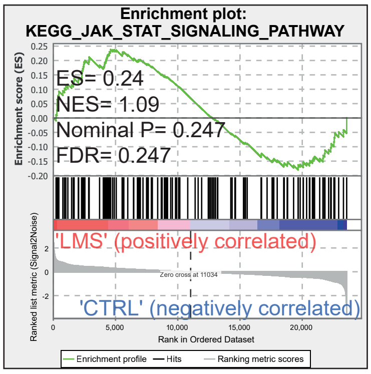


E

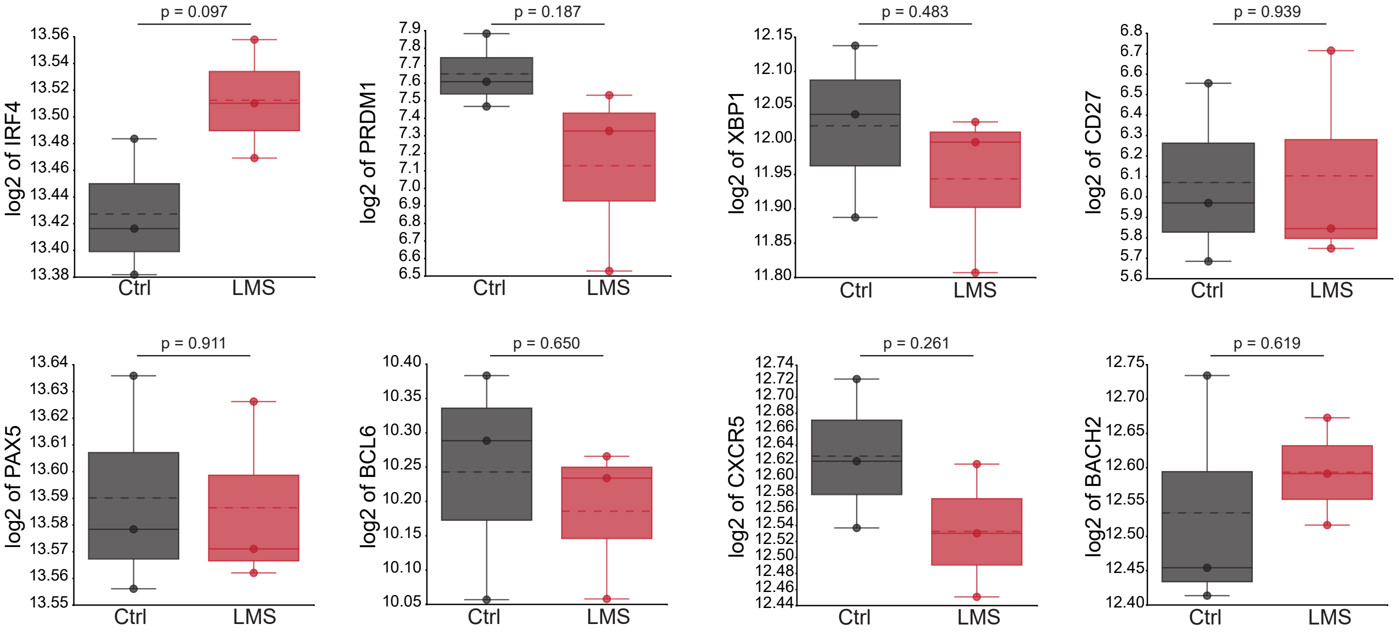

### Supplemental figure legends

#### Supplemental figure 1

LMS affects proliferation in human B-cells without inducing cytotoxicity. (A) Dot plots depicting the general gating strategy from left to right. (B and C) Titration of LMS in T-cell independent activated human B-cells in µM. B-cells were activated with CpG (5 µg/mL) for 72 h. Histograms of CFSE fluorescence (left) and quantification of percentage dividing cells gated outside parent CFSE peak (P0). (D and E) Titration of LMS in T-cell dependent activated human B-cells in µM. B-cells were activated with 3T3-CD40L cells 10:1 in combination with cytokines for 72 h. Histograms of CFSE fluorescence (left) and quantification of percentage dividing cells gated outside parent CFSE peak (P0). LMS 500 µM was chosen for consecutive experiments.

#### Supplemental figure 2

LMS causes a decrease in mitochondrial respiration but does not affect glycolysis in human B-cells. (A) Seahorse Real-Time Cell Metabolic Analysis was performed on Human B-cells from 3 healthy donors. B-cells were activated with 3T3-CD40L cells 10:1 in combination with cytokines for 48 h and treated with LMS 500 µM or solvent control. Each condition was measured in six technical replicates using a Seahorse XFe96 Analyzer to assess mitochondrial function, represented as the Oxygen Consumption Rate (OCR) over time with sequential addition of metabolic inhibitors. (B) The Extracellular Acidification Rate (ECAR) was measured in the same samples to evaluate glycolytic activity, no differences were seen between the two groups. (C) Quantification of the OCR data, shown here from left to right: Non mitochondrial respiration, Basal respiration, Maximal respiration, Proton leak, ATP production and Spare respiratory capacity. No significant differences were found (n = 3 donors, ns $P \geq 0.05$ , paired t-test).

#### Supplemental figure 3

LMS affects genes involved in cell-cycle regulation and immunoglobulin complex formation but not genes involved in DNA damage response pathways or P53 signaling. (A) Table showing leading edge genes for the gene sets HALLMARK_G2M_CHECKPOINT, HALLMARK_E2F_TARGETS and GOCC_IMMUNOGLOBULIN_COMPLEX as shown in figure 2. (B-C) ) GSEA enrichment plots show no significant enrichment of genes involved in DNA damage response or P53 signaling. Shown are WP_DNA_DAMAGE_RESPONSE, GOBP_MITOTIC_INTRA_S_DNA_DAMAGE_CHECKPOINT_SIGNALING, KEGG_P53_SIGNALING_PATHWAY and HALLMARK_P53_PATHWAY. The false discovery rate (FDR), normalized enrichment score (NES) and p-value are shown in the individual plots.

#### Supplemental figure 4

LMS treatment affects genes involved in plasma cell differentiation but not genes involved in the PI3K/AKT or JAK/STAT signaling pathways or transcription of ALPL. (A) GSEA enrichment plots show significant depletion of genes involved in plasma cell differentiation. Shown are GOBP_PLASMA_CELL_DIFFERENTIATION (left) and genes involved in plasma cell differentiation as defined by Göbel et al (Blood, 2025). The false discovery rate (FDR), normalized enrichment score (NES) and p-value are shown in the individual plots. (B) Box plots showing the transcription defined as Log2 values of ALPL in B-cells activated with 3T3-CD40L for 48 h and treated with LMS 500 µM or solvent control. No significant differences were found ($n = 3$, One-Way ANOVA with Šidáks multiple comparison test, $P = 0.936$). (C and D) GSEA enrichment plots show no significant enrichment of genes involved in the PI3K/AKT (C) and JAK/STAT (D) pathway. Shown are PID_PI3KCL_AKT_PATHWAY and KEGG_JAK_STAT_SIGNALING_PATHWAY. The false discovery rate (FDR), normalized enrichment score (NES) and p-value are shown in the individual plots. (E) Box plots showing the differences in transcription of genes involved in terminal B-cell differentiation. Shown from left to right are: IRF4, PRDM1, XBP1, CD27, PAX5, BCL6, CXCR5 and BACH2. No significant differences were found ($n = 3$, One-Way ANOVA with Šidáks multiple comparison test).
